# *In-vivo* Efficacy of a Phage Cocktail Therapy that Targets ESBL-producing *Klebsiella* species that cause Urinary Tract Infections

**DOI:** 10.1101/2025.04.01.646603

**Authors:** Rizka O.A. Jariah, Jennifer Schofield, Kevin West, Slawomir Michniewski, Eleanor Jameson, Lucy Onions, Anusha Saravanan, Hasan Yesilkaya, Andrew D. Millard, Martha R.J. Clokie, Melissa E.K. Haines

## Abstract

Uropathogenic *Klebsiella pneumoniae* strains, are of major concern of with respect to antimicrobial resistance, which has sparked interest in phage therapy as an alternative, or compliment to antibiotics. However, very limited *in vivo* data on phage therapy safety and efficacy for treating UTIs is available. To address this, we developed a model to test these parameters for a phage cocktail optimised for Extended Spectrum Beta-Lactamases (ESBL)-producing *Klebsiella* UTI strains. Female C57BL/6J mice were infected with *K. pneumoniae* Top52 then received a single dose of the purified therapeutic phages, with observations over seven days. Results showed a significant reduction of bacterial burden in urine, bladder, and kidney samples starting from 4 h post-treatment until Day 7. Phage treatment also reduced local inflammation within the bladder. Additionally, cytokine profile demonstrated an increase in anti-inflammatory IL-10 and a decrease in pro-inflammatory IL-6, indicating a synergy between phage and the host immune response.

## Introduction

Urinary tract infections (UTIs) are among the most prevalent bacterial infections worldwide, occurring in both the community and healthcare settings (1, 2). They can affect men and women of all ages, but due to anatomical differences in the genitourinary tract, infection is more common in women (3). A recent global review estimated that in 2019, over 404.6 million individuals experienced urinary tract infections (UTIs), contributing to approximately 230,000 deaths (4) alongside substantial economic and healthcare costs (2). The World Health Organization (WHO) highlighted uropathogenic bacteria including *Escherichia coli, Klebsiella pneumoniae, and Pseudomonas aeruginosa* as being major contributors to the growing global antimicrobial resistance problem (5-7). The emergence of multidrug resistance in uropathogenic strains is primarily driven by the overuse and frequent administration of antibiotics, often in the absence of proper diagnostic confirmation (8). Extended-spectrum beta-lactamases (ESBLs) represent a key resistance mechanism in Gram-negative bacteria, significantly impacting clinical management by delaying effective treatment and increasing morbidity (4, 9). Urinary tract infections (UTIs) are classified based on clinical presentation as complicated or uncomplicated. Uncomplicated UTIs occur in otherwise healthy individuals with no structural or functional abnormalities of the urinary tract, whereas complicated UTIs are associated with factors that increase the risk of treatment failure, such as urinary obstruction, catheterisation, or immunosuppression. Additionally, UTIs are categorised by infection site as lower UTIs (cystitis), affecting the bladder, or upper UTIs (pyelonephritis), involving the kidneys (10, 11). Several risk factors have been associated with UTIs, including being female, sexual activity, vaginal infection, prior UTIs, genetic susceptibility and diabetes (3, 12). In terms of the bacterial origin, it is thought that uropathogenic bacteria that can colonise the urinary tract to cause infection originate in, and are transmitted from, the normal human gut microbiota (13). In healthy individuals, these gut derived bacteria are cleared rapidly (3). Nosocomial UTIs are also common either due to catheter-associated infections or transmission outbreaks that occur between patients in hospitals (3, 14).

A major complication associated with UTIs is recurrent infections. Typically, bacteria that cause the recurrent infection are the same strain and originate from the same foci as the initial infection (15). Many different bacterial virulence factors can contribute to the recurrence of UTIs (rUTIs), such as flagella/pili, adhesins, extracellular polysaccharides, lipopolysaccharides, toxins, ureases, and proteases (9). These virulence factors allow uropathogenic bacteria to survive for long periods in a nutrient-limited habitat. It helps them to adhere, colonise, and invade host cells and to evade the host defences, then lead to persistence in the urinary tract. For example, a common uropathogenic strategy is biofilm formation, either directly on the urothelial surface or on indwelling devices such as catheters (1, 16). In rUTIs, biofilms provide uropathogenic tolerance to external stresses, including antibiotic treatments and the host immune response (1, 17).

*K. pneumoniae* is the second most abundant cause of UTIs, after *E. coli* (3, 18). There is an increased prevalence of UTI cases caused by *Klebsiella pneumoniae* producing extended-spectrum β-lactamases (ESBL); the most frequent acquired resistance mechanism in *Klebsiella* (14, 19). Medical treatment of ESBL-producing *K. pneumoniae* is therefore becoming increasingly difficult. Lack of rapid and reliable diagnostic tests to identify beta-lactamase resistance has led to the missed or late drug prescription, which eventually contributes to the widespread resistance phenotype (20) (21). The development of novel treatment strategies is clearly needed, and one potent and promising alternative is to use bacteriophages or phages, which are natural bacterial predators.

Phage therapy, where phages are used as bactericidal agents to treat bacterial infections in humans (22-24) has been used to treat infections since their discovery in 1917 (24). However, due to the success of antibiotics, phages were seen as overly complicated, and their use was largely discontinued apart from in country such as Georgia (25). With current multidrug resistance issues, phage therapy is regaining popularity, as phages have multiple attributes that make them very good antimicrobials. The first of these is that phages can replicate within their bacterial hosts to generate new phages that then replicate in other susceptible bacteria cells. This process goes on for as long the host-pathogen is present, within an infection context, this is known as self-dosing (7). Although some phages can undergo a ‘temperate life cycle’ in which their genome is integrated into the bacterial genome and replicate during bacterial cell division, their use is not recommended for therapy. This is because they can be vectors of horizontal gene transfer, and risk of spreading virulence factors, potentially making infection worse (10).

To address UTIs, *in vivo* research, *in vitro* research and clinical trials have been conducted to assess the use of phage therapy (26, 27). The most recently reported UTI phage therapy trial targeted uncomplicated UTIs caused by *E. coli* (28). The trial was deemed to be unsuccessful, but this was largely attributed to the target group actually getting better and returning to work so not completing the trial (28). In the UK, recently, there was an increase of compassionate care cases of phage therapy for cystic fibrosis patients and diabetic foot infection patients (25). However, to date, there has been only one clinical trial of phage for chronic otitis in 2009 (29).

*In vivo* models are crucial for evaluating the effects of phage therapy before its application in clinical trials. Factors such as the urinary environment, immune response, and uropathogenic characteristics impact phage efficacy and pharmacodynamics are key considerations (30, 31). Nonetheless, it is understudied in the context of UTIs (32). Recent published research assessed phage therapy efficacy *in vivo* for *Klebsiella pneumoniae* UTIs mainly through bacterial detection in urine samples (33). However, these results would not give information about the effects on intracellular bacterial communities (IBCs) or the pathology within the urinary tract. This study aimed to assess the efficacy of phages to reduce the bacterial burden of *Klebsiella pneumoniae* UTIs *in vivo* using our robust model. To do this, urine samples and and tissue samples from *the* urinary tract organs (bladder and kidney) were tested for bacterial and phage number throughout a 7-day trial period. The efficacy and safety profile of phages was determined by observing the histopathologic changes to the bladder and *by* assessing inflammatory cytokines IL-6, IFN-gamma, and TNF-alpha and IL-10 that are known to drive the resolution of infection.

## Results

### Addition of four-phages (Centimanus, Storm, Anoxic, and SoFaint) increases the *in vitro* killing efficacy against ESBL-producing clinical UTI *Klebsiella* Strains

Previously, we developed the phage cocktail-1, consisting of phages 311F and 05F, against 19 ESBL-producing *Klebsiella* strains (34). When tested against a larger set of clinically relevant ESBL-producing *Klebsiella* strains, it had a killing efficacy of 38.71% (12/31 isolates) (Figure 1A). To improve this cocktail to target a wider range of strains, we selected and tested the addition of phages that could target strains resistant to killing by phages 311F and 05F alone or in combination with each other. By the addition of phages Centimanus, Storm, Anoxic, and SoFaint to Cocktail-1, to form Cocktail-2, the efficacy was improved to ∼ 77.42% (24/31 isolates) (Figure 1B). Cocktail-2 was therefore endotoxin purified (Supp. Data 1) and utilised in further experiments.

**Figure 1.**
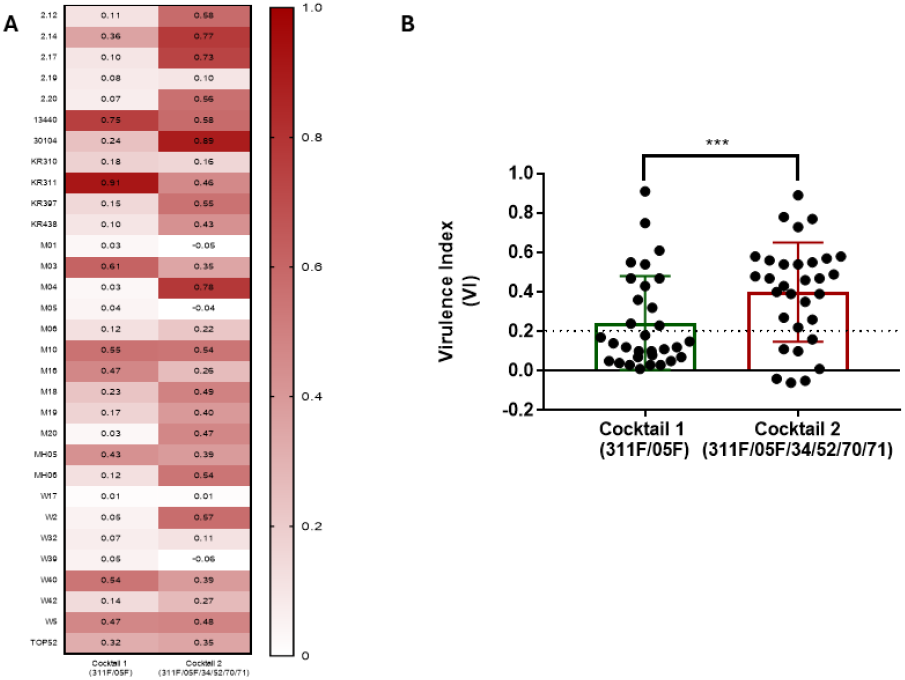
Killing efficacy comparison of Coktail-1 and Cocktail-2. a) A heat-map of local virulence index (vi) data shows the result of Cocktail-1 (311F and 05F) and Cocktail-2 (05F, 311F, 34, 52, 70, 71) against 31 ESBL-producing Klebsiella clinical isolates and K. pneumoniae Top52. The later has a better killing efficacy against tested strains b) A comparison graph showing effective killing which was defined as having VI ≥ 0.20 Cocktail 1 and Cocktail 2 to had 38.71% and 77.42% effective killing across the clinical isolate collection, respectively. ***significance difference between Cocktail-1 and Cocktail-2 (p < 0.05).

### Efficacy of transurethral phage therapy against *K. pneumoniae* Top52 UTI

We established a UTI model in C57BL/6J mice, via the transurethral route (35) using *K. pneumoniae* Top52 (Figure S1). In this study, the mice were divided into four groups: KLEB (*Klebsiella* infection control) group, KLEB-PHAGE (*Klebsiella* and phage cocktail treatment) group, PHAGE (phage cocktail only) group and CONTROL (PBS and SM-Buffer control) group (Figure 2A). We established that a starting bacterial dose of 10^7^ CFU in 50 μl would result in consistent infection and therefore 10^4^ – 10^6^ CFU/ml bacterial load could be counted until Day 7 in the homogenised bladder and in the urine. After 4 h post the bacterial or PBS injection, 5 (KLEB); 7 (KLEB-PHAGE); 5 (PHAGE); 3 (CONTROL) mice were sacrificed to determine the bacterial load in the organs (bladder and kidneys) to ensure administration was successful. Cocktail-2 was administered 48 h (Day 2) post-infection at 10^6^ PFU in 50 μl (multiplicity of infection (MOI) of 0.1) for the KLEB-PHAGE and PHAGE groups. After 4 h, 5 (KLEB); 8 (KLEB-PHAGE), 6 (PHAGE); 3 (CONTORL) of the mice were sacrificed and the rest were monitored until Day 7. For every time point, the main measured outcome was the average bacterial burden from the bladder and kidneys. From Day 0 – Day 7, the bacterial burden in the urine output was also monitored daily as non-terminal measure of infection.

**Figure 2.**
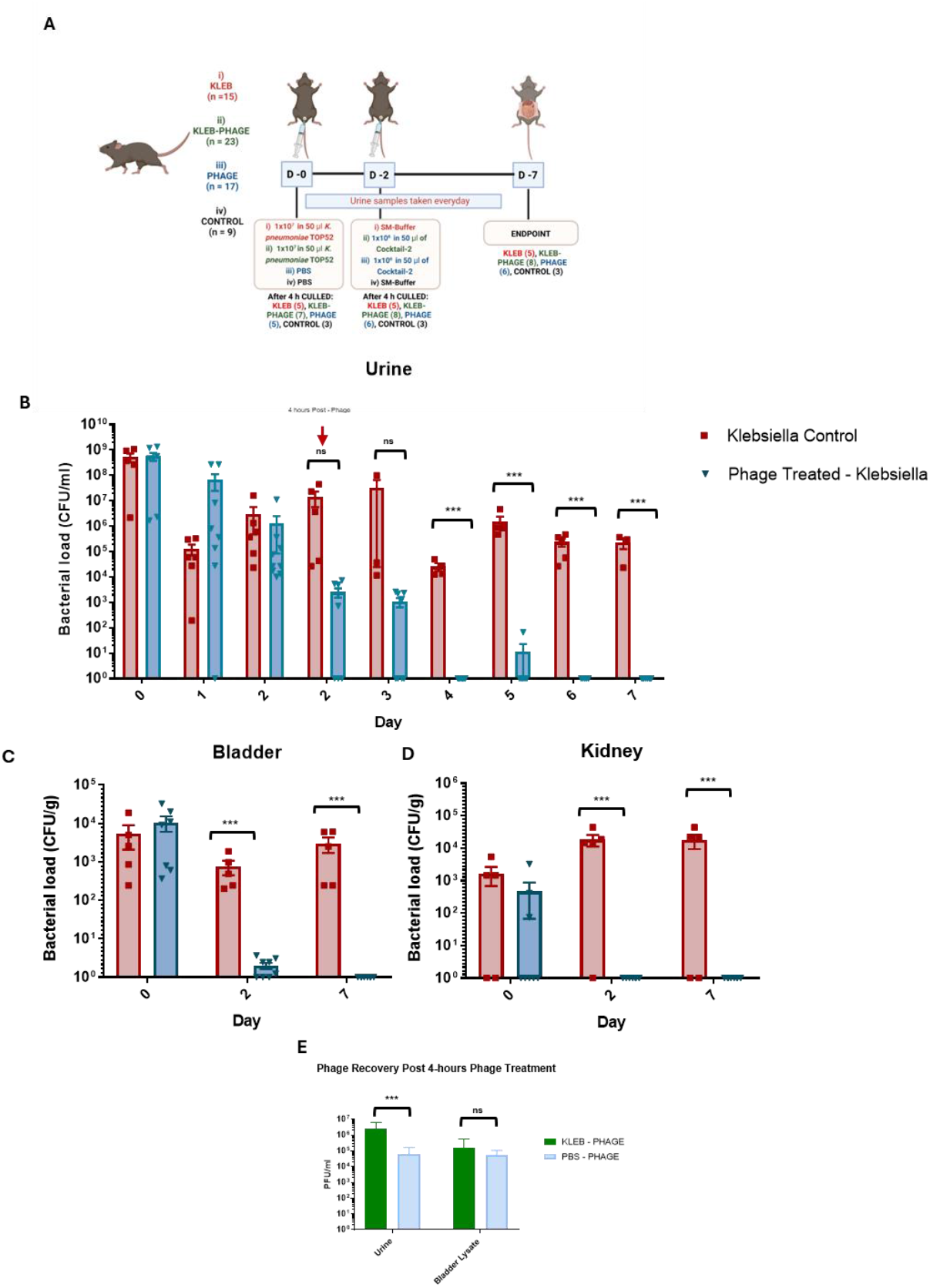
Phage therapy efficacy in a murine model. A) A murine urinary model to evaluate phage therapy against clinical strain K. pneumoniae Top52 B) bacterial burden in urine from Day 0 – Day shows significant and consistent bacterial reduction in phage treated group compared to the control; C) bacterial burden in bladder homogenates Days 0, 2 and 7 shows consistent clearance in for phage treated group; and D) bacterial burden in kidney homogenates Days 0, 2 and 7 shows total clearance of Top52 in phage treated group. E) phage recovery 4h after phage cocktail treatment on Day 2. (***p < 0.05).

*Klebsiella* levels in urine were monitored in all four groups (Figure 2B). As expected, no bacteria were detected in the PHAGE and CONTROL groups. Where an infection had been established and no phage treatment was applied, bacterial abundance ranged from 10^6^ - 10^9^ CFU/ml on Day 0 and ∼10^4^-10^6^ CFU/ml by Day 2, remaining above 10^4^ CFU/ml throughout the course of infection. Administration of phage on Day 2, resulted in an immediate drop in *Klebsiella* levels within 4 h with significantly less *Klebsiella* in urine for the phage treated mice on Days 3 – 5 and no detectable *Klebsiella* on Days 6 and 7. Thus, this shows that just one application of phage treatment appeared to eliminate the infection within the mouse bladder.

The bladder and kidney homogenates were assayed on Days 0, 2 and 7 for *Klebsiella* levels (Figure 2C and 2D). Without phage treatment, *Klebsiella* was present at ∼10^4^ CFU/g in all bladder samples (Figure 2C) until Day 7. In the KLEB-PHAGE group the bacterial burden was reduced gradually from 10^2^ CFU/g to 10^1^ CFU/g, on Day 2 and Day 7, respectively (Figure 2C). In the kidneys, *Klebsiella* was found from Day 0 of infection (4 h after bacteria injection) at 10^3^ CFU/g in the KLEB-PHAGE cohort (Figure 2D), but after Day 2, *Klebsiella* levels dropped in the phage treated group. In contrast, the levels were maintained at 10^3^-10^4^CFU/g in KLEB group throughout the time course. No *Klebsiella* was observed in the blood samples or spleen lysates for any of the mice groups, showing that no systemic infection occurred during the time course.

Phage levels were assessed after 4 h of phage transurethral injection into the bladder (Figure 2E). There were significant differences in phage recovered between the PHAGE treated group and the KLEB-PHAGE group in the urine, with 10^4^ PFU/ml and 10^6^ PFU/ml respectively. This implies the phages propagate in the urine where bacteria are present. In the bladder, both groups exhibited phage concentration of 10^5^ – 10^6^ PFU/ml at the post 4-hour treatment with no significance difference observed. No phage was detected in the KLEB or CONTROL groups as expected.

### Recovered *K. pneumoniae* Top52 colonies are still susceptible to the phage cocktail

To determine if resistance to the phage cocktail occurs i*n vivo*, bacteria were isolated from two mouse bladders on Day 7. Three isolates were randomly selected and challenged with the phage cocktail to quantify their sensitivity levels using a local virulence index (Figure 3). The three isolates 191, 197, and 200 maintained the same susceptibility as Top52 to the phage cocktail, with local virulence index scores of: 0.30, 0.32, 0.34 and 0.34 respectively. Additionally, there were no significant differences in the growth rates and doubling time of the recovered isolates compared to the original Top52 strain (Table S2).

**Figure 3.**
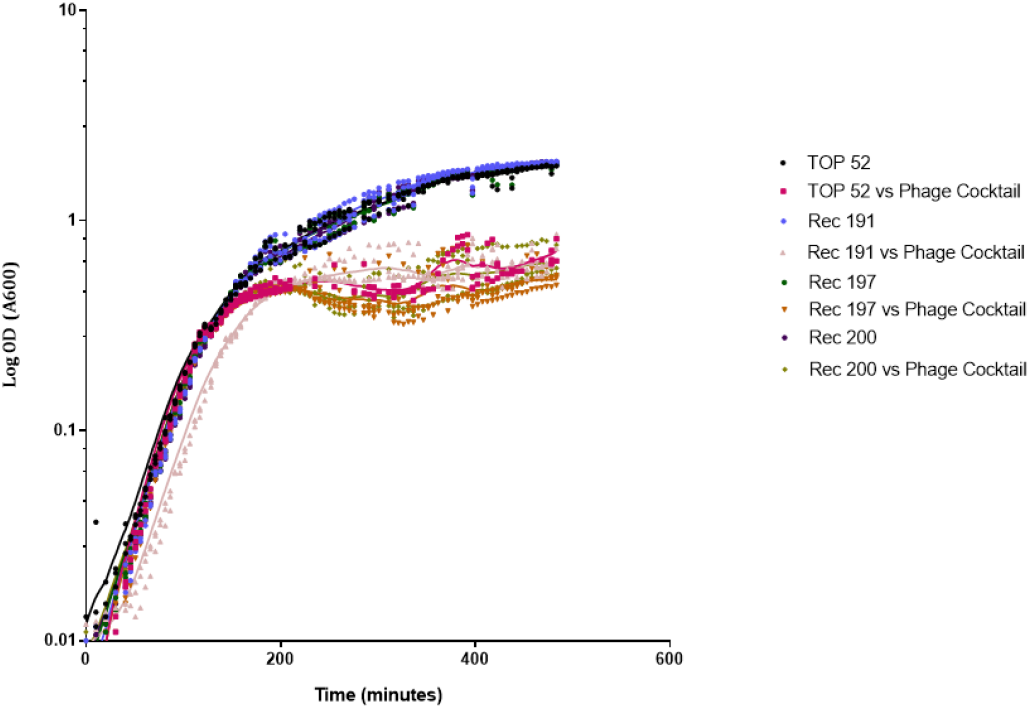
Growth curve from recovered K. pneumoniae Top52 colonies (Original Top52, Rec 191, Rec 197 and Rec 200) then challenged with Cocktail 2. OD600 was presented in a logarithmic scale with three technical replicates.

### Phage therapy provides protection for bladder tissue from severe inflammation

Histopathology analysis was used to determine the extent of inflammation that occurred during infection and the impact of phage treatment. The CONTROL mice were used as a baseline to compare all other groups to, and as expected this group had an inflammation score of zero at all time points (Figure 4A). At 4 h post-infection (Day 0), the inflammation score ranged from 1 to 4 for groups (KLEB & KLEB-PHAGE). After 4 h post phage treatment (Day 2), the KLEB group had a severe inflammation score of 4, whereas the KLEB-PHAGE group had a less severe score of 3. On Day 7, the difference was even more marked with the KLEB-PHAGE and KLEB only group having a mean score of 2.3 and 4.7 respectively. This difference in inflammation correlates with the reduced urinary bacterial burden in the KLEB-phage group (Figure 4B).

**Figure 4.**
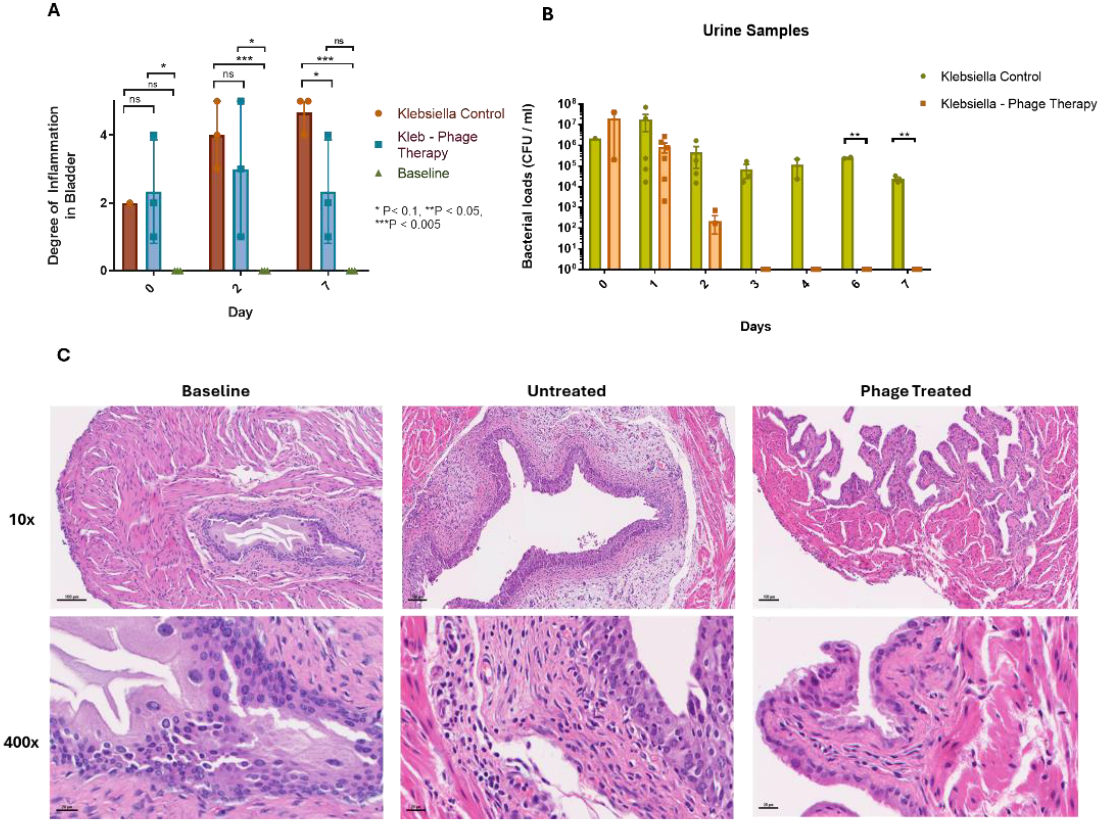
Phage therapy provides tissue protection from inflammation. A) Degree of inflammation mean score in the bladder on Days 0, 2 (4 h post phage administration), and 7 (**P<0.05) B) Bacterial loads in the urine samples taken from histology mice group Day 0 until Day 7 C) Hematoxylin and eosin staining of bladder tissue at 10x and 400x magnification (top – down respectively). From left to right, baseline inflammation score: 0; Untreated mice bladder Day 7 inflammation score: 5; Phage treated mice bladder Day 7 (post 5-days of phage treatment) inflammation score: 1.

### Pro-inflammatory (IL-6, IFN-gamma, and TNF-alpha) and anti-inflammatory (IL-10) cytokine secretion from serum and bladder lysate

To determine the impact of the phage cocktail on the inflammatory response, serum and bladder lysate samples were used to assess the expression of pro-inflammatory cytokines (IL-6, IFN-gamma, and TNF-alpha) and anti-inflammatory cytokine (IL-10). From the serum samples (Figure 5A), it was observed that the level of all pro-inflammatory cytokines: IL-6, TNF-alpha and IFN-gamma, were all elevated 4 h post *K*.*pneumoniae* Top52 inoculation. On Day 2, the IL-6 and IFN-gamma levels from the KLEB group were significantly higher than the KLEB-PHAGE group (Figure 5A, *p* < 0.05). The level of IL-6 remained high in the KLEB group continuing through to Day 7, however differing result of TNF-alpha and IFN-gamma were observed. The elevated IFN-gamma expression was also observed in PHAGE group which received PBS at Day 0, this was likely an outcome of the catheterisation process. Interestingly, anti-inflammatory IL-10 expression in serum was not observed on Day 0 in any group but was significantly higher on Day 2 in groups that had phage (KLEB-PHAGE group and PHAGE group). The level of IL-10 in the phage treated group remained high until Day 7.

**Figure 5.**
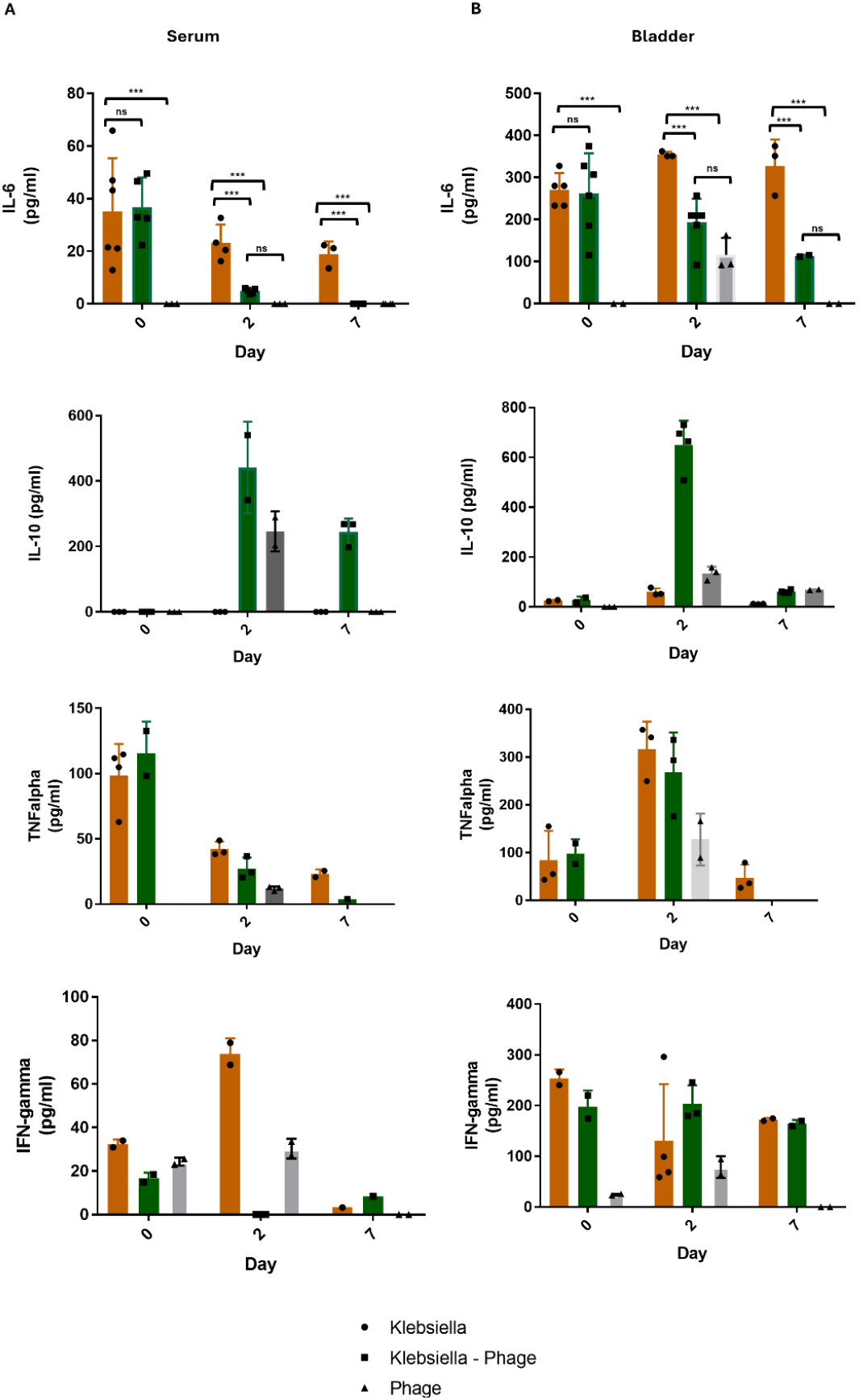
The cytokine expression to phage therapy. A) Expression of IL-6, IL-10, TNF-alpha and IFN-gamma (top-down) in pg/ml from serum samples on Day 0 (4-hours post infection), Day 2 (48-hours post infection or PBS control and 4-hours post phage treatment or SM-Buffer, Day 7. B) Expression of IL-6, IL-10, TNF-alpha and IFN-gamma (top-down) from bladder lysate samples on Day 0 (4-hours post infection), Day 2 (48-hours post infection or PBS control and 4-hours post phage treatment or SM-Buffer, Day 7. (***p<0.05)

Similar results were observed from the bladder lysate, although in general, cytokine concentrations were higher than that found in serum (10 to 100-fold). The IL-6 expression was significantly higher in KLEB group compared to the KLEB-PHAGE group (Figure 5B). However, as observed in the serum, the level of TNF-alpha and IFN-gamma level between KLEB or KLEB-PHAGE groups was not significantly different. IL-10 expression in the bladder lysate had a similar pattern to the serum, with KLEB-PHAGE group had significantly higher IL-10 expression compared to the KLEB group. In the bladder, IFN-gamma expression was constantly elevated in both KLEB and KLEB-PHAGE group (Figure 5B).

## Discussion

Although many *Klebsiella* phages have been isolated and characterised (34, 36), their clinical translation to combat *Klebsiella* UTIs is remarkably underexplored, despite *Klebsiella* being identified as the second leading cause of UTIs after *E. coli* (30) (37). Surprisingly, very few studies have evaluated the efficacy of *K. pneumoniae* phage therapy *in vivo*, or in animal models (33, 38). Indeed, no previous *Klebsiella* UTIs mouse models have been developed to study phage efficacy. To address this our study aimed to bridge this gap by developing a robust *in vivo* model in which to assess the efficacy and safety of a phage cocktail in treating *Klebsiella*-associated UTIs. To do this, an established *Klebsiella* UTI *in vivo* model was established in our pre-clinical facility and evaluated to determine the safety and efficacy of phage therapy to reduce bacterial disease as assessed by a reduction of bacterial numbers and physical and immune responses to the phages. As such, in addition to this microbiology data, to determine how phages interact with a mammalian immune system we examined infection-induced inflammation using histopathological analysis and cytokine expression profiling

In this study, previously developed phage cocktail in our laboratory that targets ESBL-producing clinical UTI *Klebsiella* isolates was further developed to ensure clinical relevance in terms of spectrum of activity on relevant strains, for individuals with limited antibiotic treatment options. The collection of clinical isolates was compiled from multiple locations across the UK, including Leicester, Manchester, Bristol, and Public Health Wales, to capture a representative diversity of clinically relevant and prevalent strains (Table 2). Previously published *in vitro* studies from our laboratory on an optimised phage cocktails that targeted ESBL-producing *Klebsiella* strains identified Cocktail-1 (311F and 05F) as having the broadest spectrum of activity at 53% (34). Building on this, the current study tested Cocktail-1 against a different subset of ESBL-producing *Klebsiella* strains and incorporated four additional phages to broaden its killing spectrum, called Cocktail-2 (311F, 05F, Centimanus, Storm, Anoxic, and SoFaint). The efficacy of the new combination was evaluated using a phage killing assay (PKA), and comparisons between different phage cocktails were made using the local virulence index (*vi*) score—a quantitative method that compares the area under the growth curve of bacteria following phage addition to a standard bacteria only growth curve (39). Phage cocktail activity is generally assessed using two key criteria; the first being the breadth of activity, which refers to the number of isolates the cocktail can infect, to identify those capable of targeting a wide range of strains. The second is the depth or strength of activity, which measures the ability to reduce bacterial numbers. Although we value both parameters for clinical applications it is essential to target as many as possible clinical strains and thus we have prioritised this parameter over a cocktail with a narrower spectrum (40).

Comparison of Cocktail-1 and Cocktail-2 (Figure 1A and 1B) revealed that the addition of four new phages increased the killing efficacy and expanded the cocktail’s spectrum of activity. Cocktail-2 demonstrated greater breadth of activity, targeting more strains than Cocktail-1. While the local virulence index (*vi*) for some strains was classified as medium (*vi* 0.20 - 0.50), the phage cocktail still effectively reduced bacterial growth. The findings suggest there is synergy between the six phages, where the growth and infection dynamics of one phage influence that of the others within the same bacterium. Synergistic interactions can affect key lytic phage characteristics, such as adsorption rate (infection efficiency), burst size (number of progeny produced), and lysis time (duration from infection to progeny release) (41). Indeed, genomic analysis (Table 3) conducted by collaborators (Eleanor Jameson *et al*.) provided additional insights, confirming that phage Centimanus in Cocktail-2 encodes nine depolymerase proteins. These enzymes disassembly bacterial capsules, facilitating phage infection (41, 42). While these findings highlight the potential of Centimanus in enhancing cocktail efficacy, further research is necessary to elucidate the detailed mechanisms of its depolymerase activity (43).

**Table 1.**
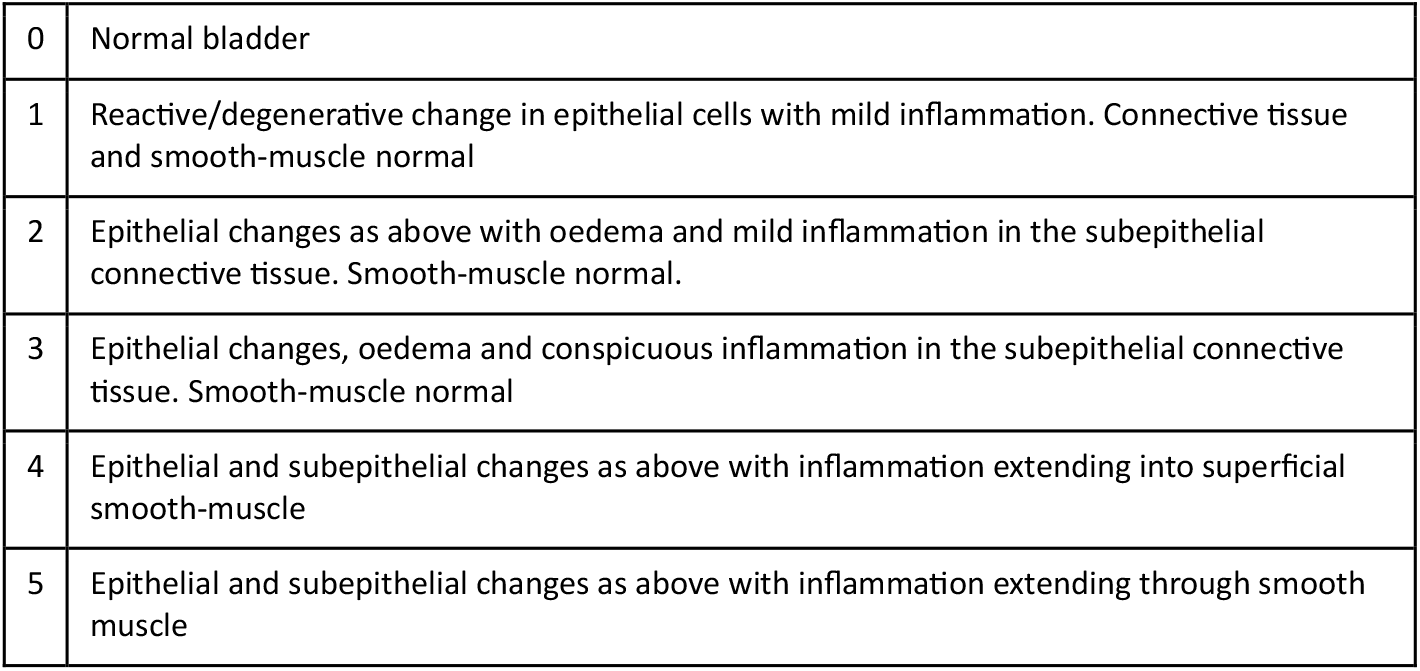
Histopathology grading scale for the degree of inflammation in the bladder tissue. Ranging from 0 – 5, with 0 representing no inflammation and 5 representing the most severe and extending inflammation.

**Table 2.**
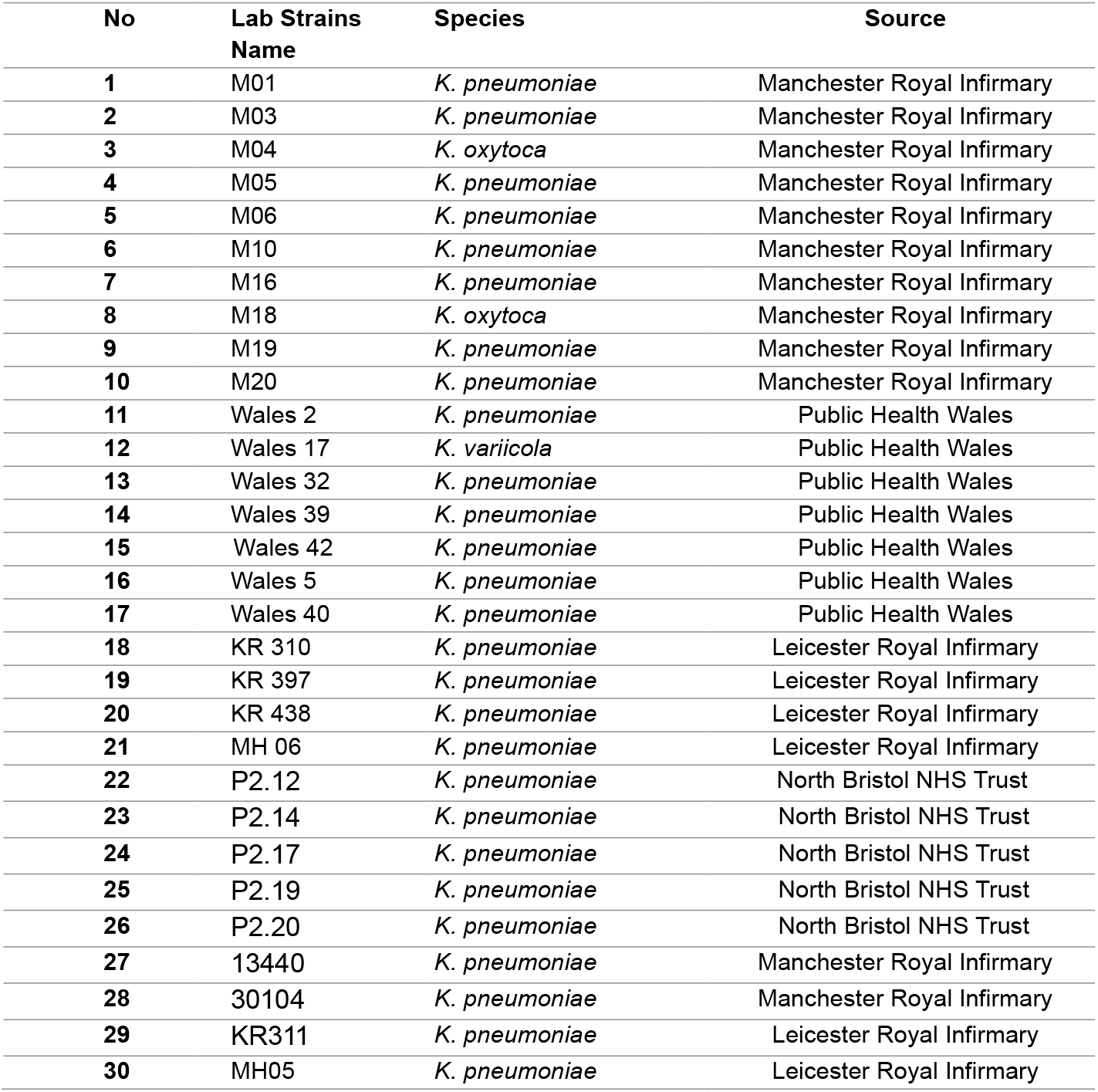
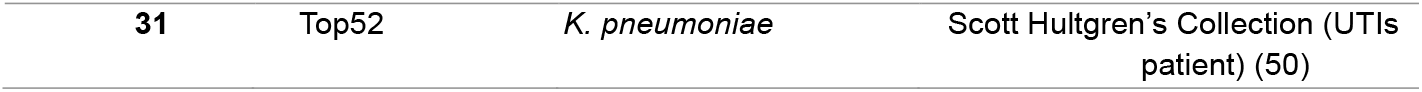
Klebsiella spp. clinical strains used in the host range study; All ESBL-producing except Top52.

**Table 3.**
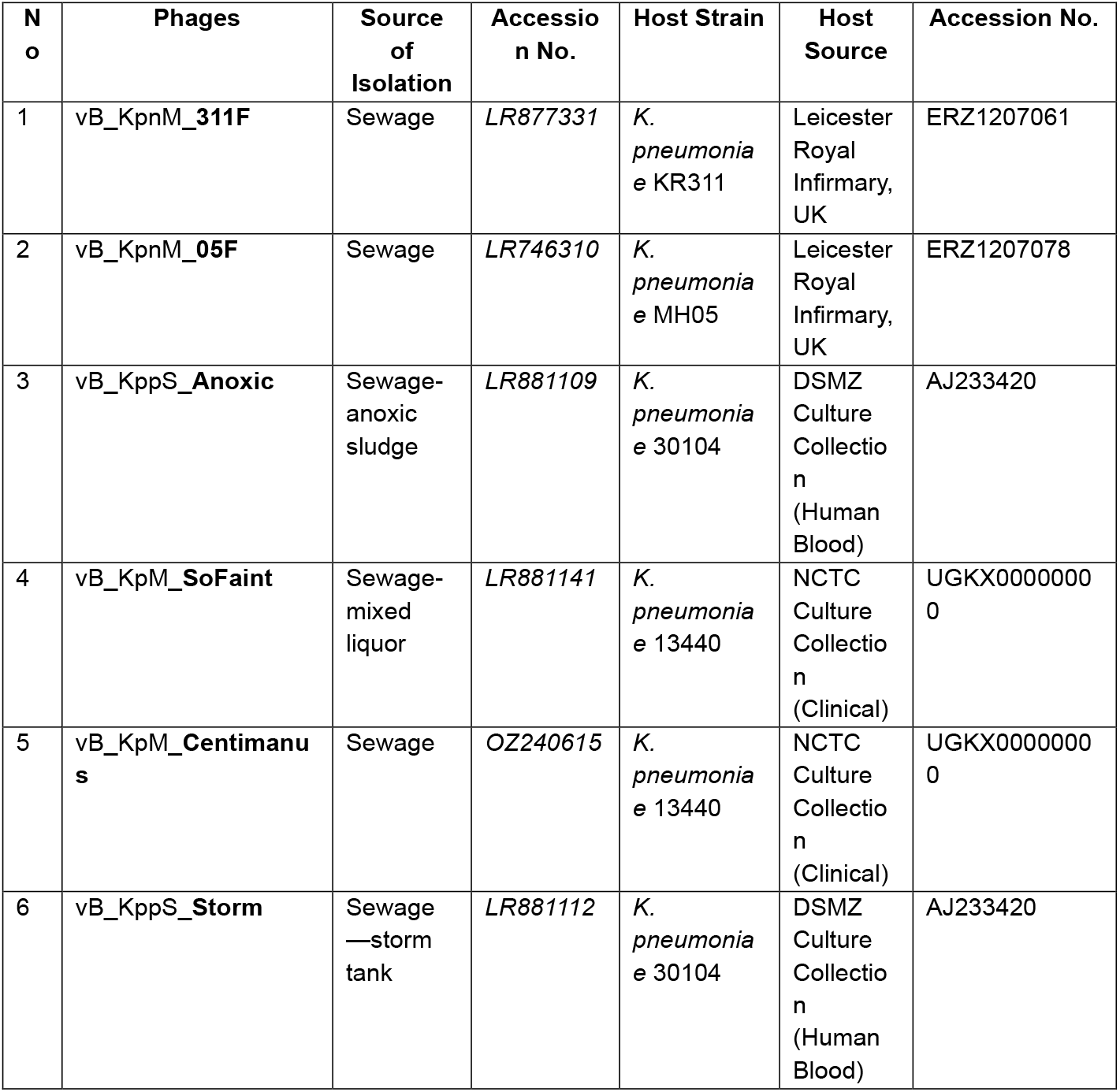
Phages and original propagation hosts used in this study.

Murine models have proven invaluable for studying UTIs in non-phage contexts but have not been previously adapted to study how phages might behave within a complex bladder environment. Unlike other rodent models, mice do not naturally exhibit vesicoureteral reflux which can lead to UTIs (44). However, they share physiological similarities with humans, such as a comparable proportion of globoseries glycolipid receptors on urothelial cells, which facilitate bacterial attachment (45), and conserved uroplakins that promote type 1 fimbriae adherence and UPEC invasion (46). While mice do not naturally develop UTIs, several instillation techniques, including unobstructed UTI instillation and transurethral catheterisation, have been developed to experimentally induce infections (47, 48). The catheterisation method is particularly advantageous to probe UTIs, as it allows precise control over infection severity so a localised or systemic infection can be induced.

In this study, we used the *K. pneumoniae* Top52 strain, which was originally isolated from a female patient with recurrent UTIs and has previously been used in murine pneumonia model (49, 50). Encouragingly, within our C57BL/6J mouse lines, persistent high bacterial titres (>104 CFU/mL) were maintained over seven days, mimicking a sustained bladder infection (66). Selecting the appropriate bacterial strain is critical to establishing murine UTI models, as phenotypic differences in bacterial behaviour can arise between human and mouse environments due to variations in development and growth conditions (67). Additionally, bacterial virulence factors also dictate behaviour, therefore the selected strains should exhibit similar pathologies that closely model the UTIs (1).

One of the problems that makes UTIs bacteria so problematic is that they can form intracellular bacterial communities (IBCs) (51). Upon invading bladder epithelial cells, these bacteria proliferate within the cytoplasm and expand into the bladder lumen as colonies grow (51). These colonies can form biofilm-like structures, invade deeper urothelial layers, and persist in membrane-bound compartments with limited metabolic activity—a state known as “quiescent intracellular reservoirs” (QIR). These reservoirs remain latent but can reactivate when the upper urothelium exfoliates (52, 53).

Our results demonstrated that although the *in vitro* virulence index (*vi*) for Cocktail-2 against Top52 was 0.35 (considered to be medium according to Haines *et al*) (34), a complete clearance of bacterial burden in the urine and bladder was achieved on study Day 7. This suggests that the phage cocktail is more effective in an in vivo system than in intro and that the phages may effectively target IBCs within the bladder. We hypothesise that the observed clearance of bacteria is a consequence of the synergy between phages and the murine immune response, and thus the phages have driven an efficient infection resolution. By reducing bacterial load, phage therapy may enhance the immune system’s ability to eradicate residual bacterial populations (54). Furthermore, phages can promote bacterial phagocytosis by macrophages, as they coat bacterial surfaces, increasing their susceptibility to immune response mechanisms (55, 56). Thus, understanding how phages interact with the immune system during therapy is a crucial aspect of developing effective treatments.

A histopathological analysis (Figures 4A and 4C) revealed that *K. pneumoniae* Top52 formed IBCs, invaded urothelial cells, and triggered inflammation extending into the smooth muscle. In contrast, phage treatment appeared to protect against severe inflammation and potentially facilitated tissue recovery. This effect may of course result from the reduced bacterial burden achieved through phage therapy, but our data suggest that the picture is more complex. Indeed, phages have been shown to elicit diverse immune responses in humans (48). For instance, Pf4 phage treatment for *Pseudomonas aeruginosa* infections has been shown to induce IFN-β and downregulate pro-inflammatory TNF (57). Upregulation of anti-inflammatory cytokines after phage application has also been observed in a murine study which showed injected phages induced systemic IFN-β, which provided additional protection against vaccinia virus infections (58). Of critical relevance to this study is that, phages have been reported to induce IL-10 expression, which is an anti-inflammatory and regulatory cytokine that mitigates against tissue damage (59, 60). The *in vitro* finding of phage application effect on monocytes is consistent with our observations (59). In our study, the mice that received phage therapy had upregulation of systemic and localize IL-10. IL-10 has been extensively studied as a major tissue-protective cytokines, which are able to protect the host against Th1 and Th2 associated tissue damage exerted by bacterial infections (60). This explains lower inflammation score in the bladder tissue of phage treated mice compared to the infection control as phages induced IL-10 and suppressed tissue inflammation. Although IL-10 production often coincides with IL-6 expression—a critical inflammatory cytokine involved in viral resolution and Th2 cell expression, which is also essential to get a balanced immune response (57, 61, 62).

While this study highlights the potential of phage therapy as a treatment for UTIs, it is important to acknowledge the study limitations. One limitation was that the volume of urine retrieved from mice could not be standardised, as not all mice produced daily urine samples. Efforts were made to address this using metabolic cages and positive reinforcement with treats, yet occasional failures to collect samples persisted. Secondly, the relatively short study duration of seven days means that understanding treatment for chronic infections lasting more than two weeks or recurrent UTIs is limited. Therefore, running longer experiment might be necessary to test phage for chronic UTIs treatment.

In summary, we showed for the first time that we have a system to explore the efficacy of phages in a robust preclinical model which has allowed us to gather important insights into how phages could be exploited to treat UTIs in humans. The robust dataset demonstrates that, for uncomplicated acute UTIs, a single dose of the phage cocktail is sufficient to remove the bacterial pathogen and resolve the infection. Furthermore, the observed anti-inflammatory response exerted in the phage-treated group through transurethral injection underscore the potential applicability of phage cocktails in reducing inflammation caused by UTIs The histopathology data also supports the anti-inflammatory effects of phage therapy, which is critical to solve the infection.

In future work the cocktail could be expanded to address polymicrobial infections, for example, incorporating additional phages to target ing other bacterial species. For instance, prior research by Haines et al. (34) combined *E. coli* and *Klebsiella* phages, demonstrating efficacy towards both species within the mixed cocktail. Additionally, future research on the use of phage-antibiotic combinations *in vivo* will be essential as phages are often used in conjunction with antibiotics rather than as standalone therapies (63). Notably, therapeutic synergy between antibiotics and phages has been shown to improve patient outcomes in cases where phages have been used under compassionate grounds (64-67).

## Materials and Methods

### Strains and Phages Used in This Study

31 *Klebsiella* strains were used in this study; 30 were ESBL-producing *Klebsiella strains*, previously isolated from patients with UTIs were used to test phage host range (Table 2). The final strain was Top52 #1721 (Top52) that was sourced from Scott J. Hulgren and used *in* the in *vivo* mouse *model* (50). All strains were cultured from frozen stocks on LB agar medium (1.5% w/v, Luria Bertani media Sigma-Aldrich, United Kingdom L3522, Bacteriological Agar, VWR, United Kingdom), supplemented with 1 mM CaCl2 (Fisher Chemical, Fisher Scientific, United Kingdom) and incubated at 37 °C overnight (∼18 h). The liquid culture was grown in LB medium and incubated at 37 °C in 300 rpm shaking incubator overnight.

Phages used in this study was sourced from the Becky Mayer Centre for Phage Research collection (Table 3).

### Phage Propagation, Titration and Purification

The double-layer agar method was used for each phage propagation from the stock culture. In brief, bacterial strains were cultured to log phase (OD600 = 0.15 – 0.2) using an overnight culture as a starting point. 200 μl of bacteria culture and 200 μl of phage stock were then added to 8 ml of 0.7% (w/v) LB agar, poured onto square LB 1.5% (w/v) agar plates and incubated 37 °C overnight (∼18 h). The next day, the top layer agar was scraped and transferred into a 50 ml Centrifuge Tube with 10 ml SM Buffer [100 mM NaCl (Sigma-Aldrich, United Kingdom), 8 mmMgSO4•7H_2_O (Sigma-Aldrich, United Kingdom), 50 mM Tris-HCl pH 7.5 (Sigma-Aldrich, United Kingdom)] and incubated at room temperature for ∼18 h. The phage lysate was centrifuged at 4,000 × g for 15 min and the supernatant was filter-sterilised through 0.2 μm pore size filters. The phage stock titre was determined using plaque assays. The phage stocks were stored at 4°C.

### Phage Host Range with Planktonic Killing Assay

To determine the host range of the cocktail, a planktonic killing assay (PKA) was performed using a previously described method (34). This optimisation step was the development of the previously described *Klebsiella* phage cocktail: 311F and 05F (Cocktail 1). We then developed further by testing the addition of phages Anoxic, SoFaint, Centimanus and Storm into the cocktail (Cocktail 2). Briefly, bacteria overnight cultures were diluted 1:100 and added to flat bottom 96 well plates in triplicate per strain. Negative controls used LB only and Blanks used LB with Gentamicin 2 μg/ml. The bacteria grown to OD600 0.15 (1 × 10^8^ CFU/ml) and 100 μl of phage cocktail (containing 10^8^ PFU/ml of each individual phage) was added in each well, and dilution of phages was made using LB media. The PKA was done at a MOI of 1. The experiments were performed using the BMG Labtech SPECTROstar Omega. The program was set up as follows: OD600 taken every 5 min for 8 h with shaking 10 s before each reading. The result was merged using the program to produce a killing assay curve. The curve was then analysed by a method developed by Storms et al (39). The local virulence index (*vi*) score was a ratio of area under the bacteria growth curve challenged with phage cocktails against the bacteria growth curve without phage (positive control) during the log phase. The local virulence index gives a value between 0 to 1, 0 means not effective and 1 means highly effective. Effective killing in this study, was considered if the virulence index was ≥ 0.20.

### Phage Endotoxin Purification

Endotoxin purification is required for phage products to fulfil the minimal standard for a clinical product. For intravenous application, the limit is 5 EU/ml and for oral administration it is < 20 EU/ml. Endotoxin purification was carried out using commercial endotoxin affinity assay Endotrap HD (LIONEX, GmBH) following the manufacturer’s instructions. In brief, 1 ml of purified phage in SM-Buffer was run through the column using gravity column. 200 μl of void volume was collected in a separate tube. The total phage lysate volume was run through was 20 ml. Endotoxin quantification level was performed using Pierce Chromogenic LAL Assay (ThermoFisher, United Kingdom) as per its instructions. The purified Cocktail-2 lysate was shown to retain the same killing ability as the unpurified version (Figure S2).

### Experiment Design of Infection Model *In vivo*

For the *in vivo* study, the method was modelled according to the well-established protocol (35) (Supp. Fig 2). *K. pneumoniae* Top52 was cultured in LB medium and incubated statically for 18–24 h at 37 °C, then sub-cultured 20 μL into 20 ml fresh LB broth and incubated statically again for 18 h at 37 °C. The culture was centrifuged at 4200 RPM for 10 min, pellet was resuspended in 10 ml sterile PBS (Sigma Aldrich, United Kingdom). The OD600 was normalized to 0.91 – 1.00 this corresponds to approximately 2 – 4 × 10^8^ CFU/ml (1 - 2 × 10^7^ CFU in 50 μL). The titre was verified using serial dilutions and spotting method on cystine-lactose-electrolyte-deficient (CLED) agar medium (VWR, No. 75813, USA)

7–8-week-old female C57Bl/6J mice were bred and maintained in-house at the University of Leicester. Mice were caged in a room with a 12/12 h light-dark cycle and had ad libitum access to water and food pellets. In the experiment, mice were randomised into 4 groups.: Group 1 KLEB (n= 15) with *K. pneumoniae* Top52 at 0 h and SM-Buffer at 48 h, Group 2 KLEB-PHAGE (n= 23) with *K. pneumoniae* Top52 at 0 h and 10^6^ PFU in a total volume of 50 μL PHAGE cocktail at 48 h, Group 3 PHAGE (n= 17) with PBS at 0 h and 10^6^ Phage cocktail at 48 h, Group 4 CONTROL (n= 9) with PBS at 0 h and SM-Buffer at 48 h (figure 2A). Mice were lightly anesthetised with 2.5% (v/v) isoflurane over oxygen (1.8 - 2 L/min) and the procedure was done via transurethral injection with 50 μL volume. At each timepoint, mice were humanely killed by terminal anaesthesia and immediately dissected. Bacterial levels at Day 0, Day 2 (4 h after interventation either phage treatment or PBS) and Day 7 were defined as the abundance of *K. pneumoniae* Top52 per ml in bladder, kidneys, and spleen tissue lysate. Urine samples were attempted to be collected from all mice every day starting from Day 1 up to Day 7. Positive reward training or bladder massaging method were used to encourage the mice to urinate on the petri dish. Alternatively, the mice were taken into metabolism cages for 1 – 3 h until they produced urine. The bladder, kidneys, and spleen were homogenised using FastPrep Homogenization (FastPrep 24). The homogenised organs were then serially diluted and plated on CLED agar plates for bacterial enumeration. The rest of the homogenised solutions were processed for phage counting by using the plaque assay method on double-layer agar as mentioned above. Blood samples were also taken via cardiac puncture during the terminal anaesthesia procedure and samples were processed for counting bacterial load. Additional mice were used for histopathology analysis following the same infection model.

### Histopathology

The bladder and kidneys of the mice were reserved for histopathology assessment in 10% formalin (VWR International, United Kingdom). Organ sections of 5 μm thick were cut from paraffin-embedded (Leica HistoCore Pearl Tissue Processor) and stained with hematoxylin and eosin. The histopathological grading scale for the degree of inflammation (0-5) was adapted from the original definition by Hopkins (39), with additional detail changes presented in Table 1. Slides from the CONTROL group (untreated mice) were used as a baseline for uninfected tissue. A blinding method was applied during the grading process, so that the treatment groups and culling time points were kept confidential to limit reporting bias.

### Phage Susceptibility Testing of *Klebsiella* colonies Post Murine Infection

*Klebsiella pneumoniae* Top52 colonies were isolated from bladder or urine samples of Group 2 KLEB-PHAGE. Colonies were streaked on CLED agar plates and incubated overnight at 37 °C. The colonies were re-streaked two times on CLED agar plates. The plates then stored at 4 °C prior to phage sensitivity testing. Recovered colonies were inoculated in LB broth and incubated overnight at 37 °C in 300 rpm shaking incubator. Sensitivity of recovered colonies were tested against Cocktail 2 (311F, 05F, Anoxic, SoFaint, Centimanus and Storm) with 10^8^ PFU/ml concentration (MOI 1) using PKA method. The local virulence index was calculated as previously explained. The growth and doubling times of recovered bacteria was analysed using DASH (68) (Supp. Fig 4).

### IL-6, IL-10, TNF-alpha and IFN-Gamma Expression

The serum was retained from mouse blood by centrifuging the fresh blood in 4°C at 9000 RPM and was stored at -20°C. The bladder lysate was kept at -20°C and centrifuged before the ELISA was performed. The ELISA was performed using commercially available kit (BioLegend, IL-6 Cat. no 431304, IL-10 Cat. no 431414, TNF-alpha Cat. no 430904, IFN-gamma Cat. no 430804) as per the instructions. All samples were done in triplicate.

### Ethics Statement

All experiments were in accordance and approved by Animal Welfare Ethical Review Board (AWERB) at The University of Leicester Ethical Committee and the U.K. Home Office under project license PP0757060.

### Statistics

Graphic and statistical analyses were performed using GraphPad Prism 7 (GraphPad Software, Inc). For both phage cocktail comparison and animal experiments with two groups, the variances of the outcome variables were compared to select the appropriate statistical tests. Shapiro-Wilk test was performed to verify normal distribution. A two-tailed t-test if the data was normally distributed, or two-tailed Mann-Whitney U-test if not. All *in vitro* experiments were performed in triplicate.

## Acknowledgements

We acknowledge the help and support from the staff of the Division of Biomedical Services, Preclinical Research Facility, University of Leicester, for technical support and the care of experimental animals. We also like to thank Leicester Core Biotechnology Histology Services for the assistance with histology staining and Advanced Imaging Facility (RRID:SCR_020967) at the University of Leicester for support with histology imaging. This work was supported through the Institute for Precision Health and Leicester Drug Discovery and Diagnostics (LD3) via the institutional MRC Impact Accelerator Account [MR/X502777/1] and Catto Foundation. Thank you for Public Health Wales, University Hospitals of Leicester, Manchester Royal Infirmary, North Bristol NHS Trust (Pippa Griffin) and Scott J. Hulgren for providing bacterial isolates collections.

R.O.A.J PhD Study was funded by Beasiswa Pendidikan Indonesia (BPI), Indonesia Endowment Fund for Education (LPDP) and Center for Higher Education Funding and Assessment (PPAPT), Ministry of Higher Education, Science, and Technology of Republic Indonesia. M.E.K.H, Academic Clinical Lecturer is funded by the NIHR. The views expressed in this publication are those of the author(s) and not necessarily those of the NIHR, NHS or the UK Department of Health and Social Care. A.M is funded by MRC (MR/L015080/1 and MR/T030062/1). Bioinformatics analysis was carried out on infrastructure provided by MRC-CLIMB (MR/L015080/1).

For the purpose of open access, the author has applied a Creative Commons Attribution license (CC BY) to any Author Accepted Manuscript version arising. Figure 2A was created in BioRender by R.O.A.J (2025) https://BioRender.com/1uc0l27

## Author Contributions

R.O.A.J, J.S, H.Y, M.R.J.C and M.E.K.H conceived the experiment, S.M and E.J performed phage isolation, R.O.A.J, A.S, J.S, and L.O performed the experiment, R.O.A.J, K.W and S.M analysed the data, R.O.A.J, A.D.M, M.R.J.C and M.E.K.H wrote the manuscript.

