## Supplementary Information for "*In-vivo* Efficacy of a Phage Cocktail Therapy that Targets ESBL-producing *Klebsiella* species that cause Urinary Tract Infections"

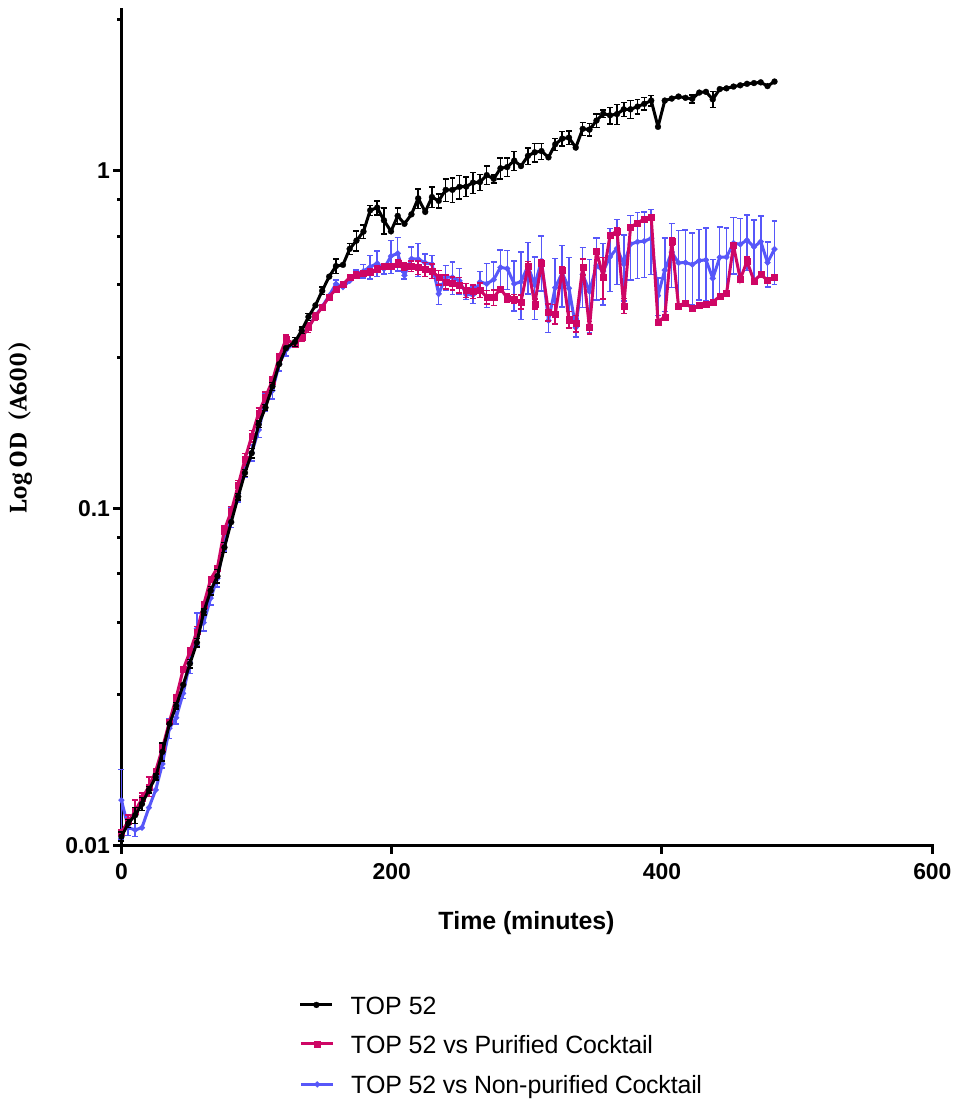

- Supplementary figure 1. Killing curve of *K.pneumoniae* Top52 against Purified or non-purified Phage cocktail at MOI 1. The purified cocktail still has the same killing effect as the non-purified. The local virulence index (*vi)* was 0.37 and 0.35, for purified and non-purified phage cocktail, respectively.

Supplementary figure 1. Killing curve of *K.pneumoniae* Top52 against Purified or non-purified Phage cocktail at MOI 1. The purified cocktail still has the same killing effect as the non-purified. The local virulence index (*vi)* was 0.37 and 0.35, for purified and non-purified phage cocktail, respectively.

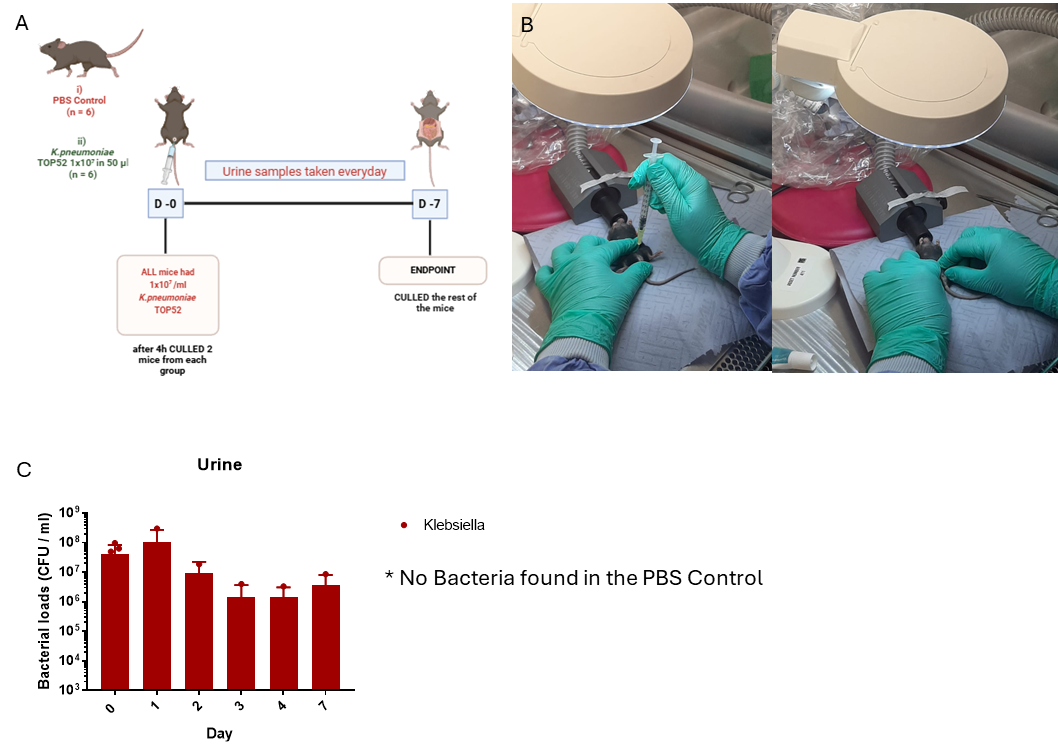

Supplementary figure 2. A) Established model for *Klebsiella-*UTIs mice model. Optimum dosage was 10^7^ CFU given in 50 µl via transurethral injection which can cause consistent infection for 7 days. B) Transurethral injection process to the mice using catheterisation method. C) Bacterial load data from urine samples, taken daily Day-0 – Day 7 showed stable infection ranging from 10^7^ – 10^6^ CFU/ml.

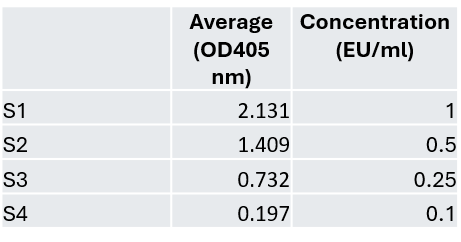

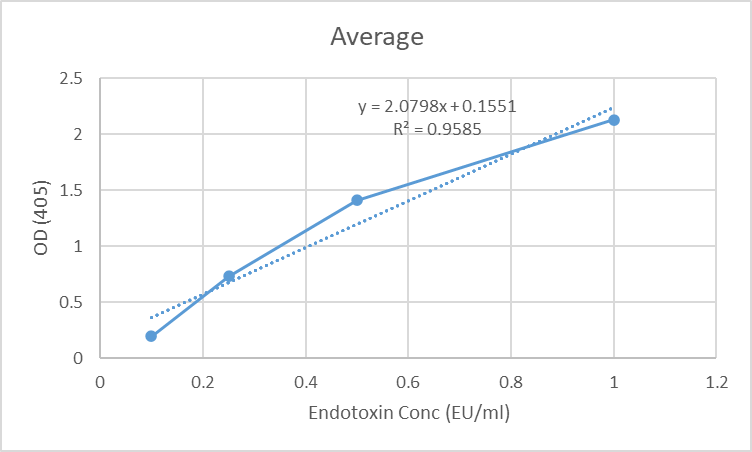

**Result:**

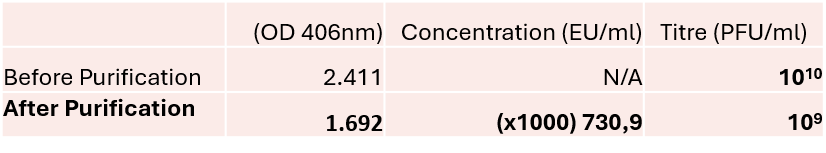

Supplementary Data 1. Endotoxin Quantification from Purified Phage Lysate

| **Phage Name**  Supplementary table 1. List of phages used in this study. | **TEM** | **Genus** | **Genome Size** | **Reference** |
| --- | --- | --- | --- | --- |
| vB_KpnM_311F | 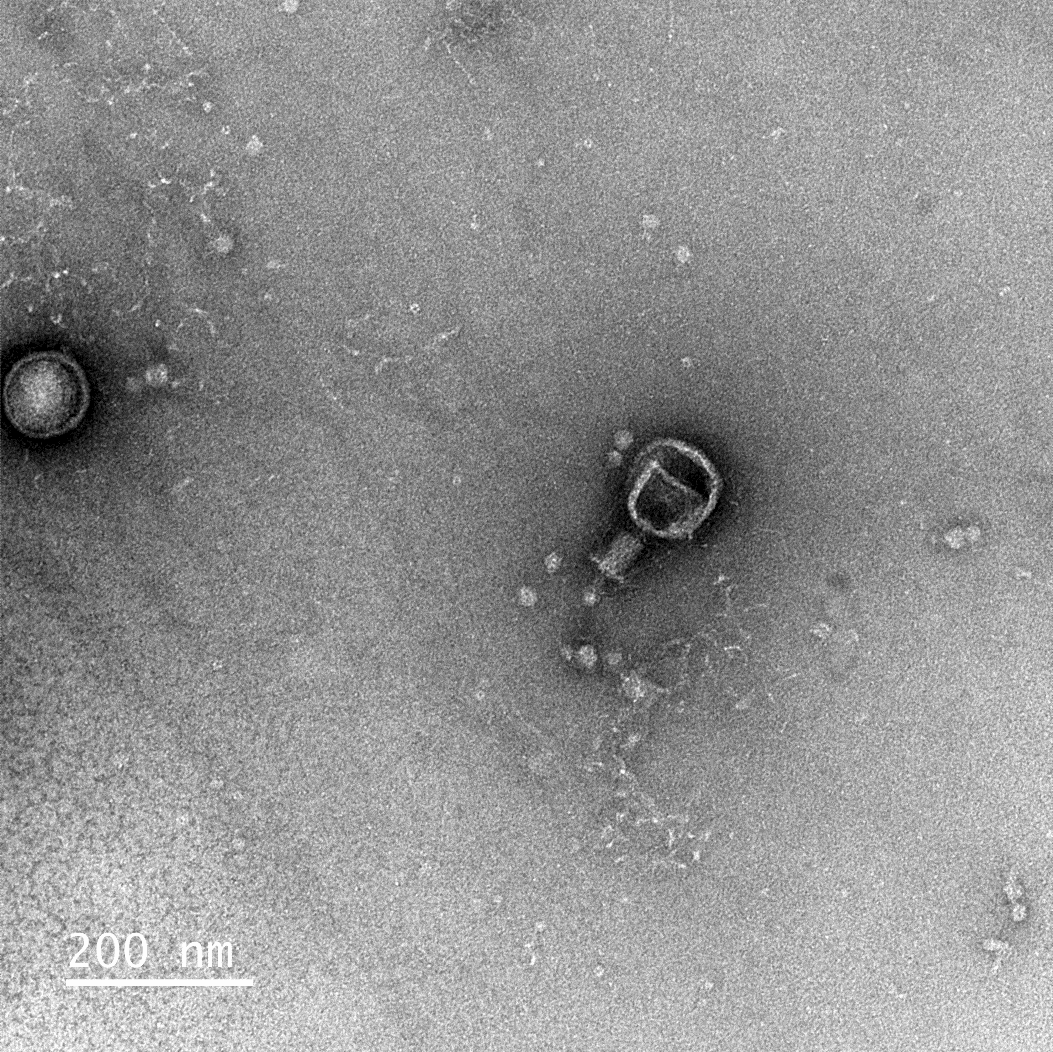 | *Jiaodavirus* | 186 kb | 34 |
| vB_KpnM_05F | 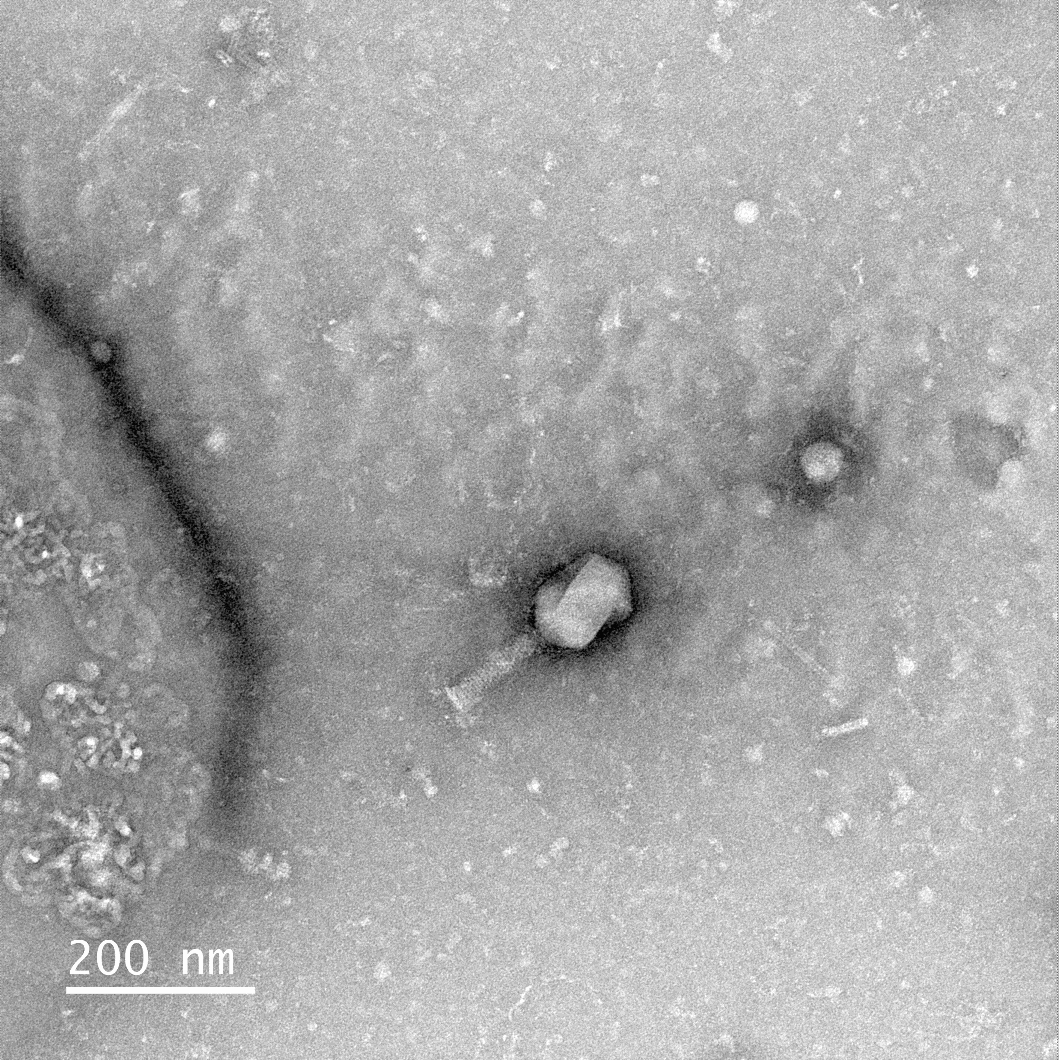 | *Jiaodavirus* | 175 kb | 34 |
| vB_KpM_Centimanus (Phage 71) | 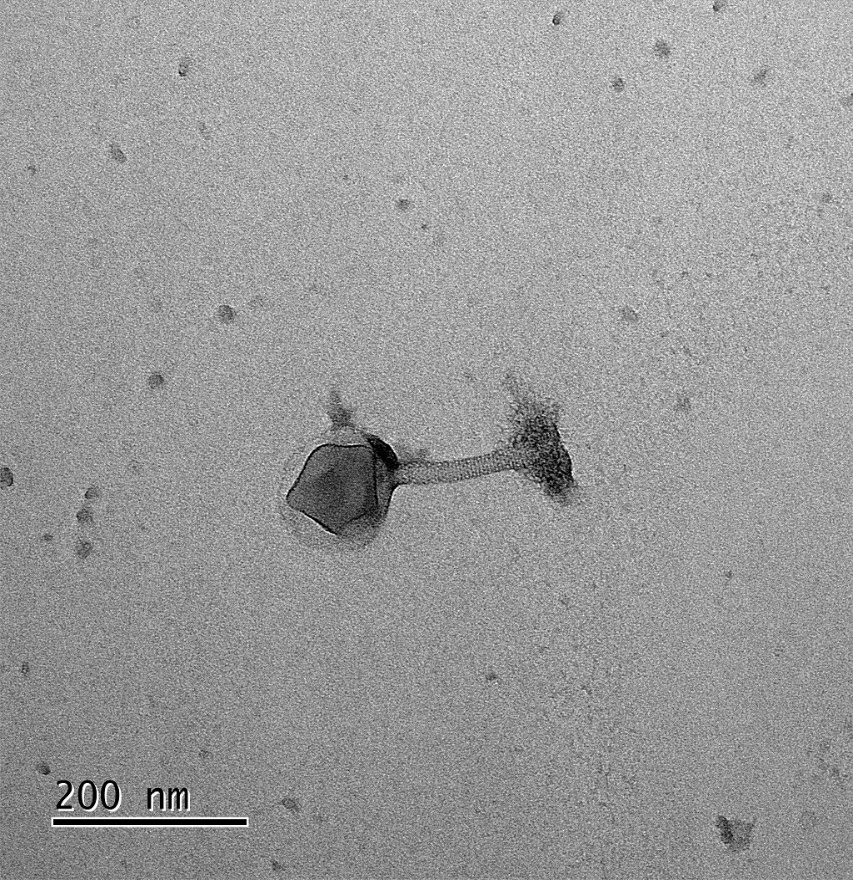 | *Maaswegvirus* | 299 kb | - |
| vB_KppS_Storm (Phage 34) | 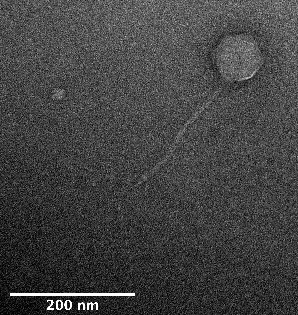 | *Sugarlandvirus* | 110 kb | 36 |
| vB_KppS_Anoxic (Phage 52) | 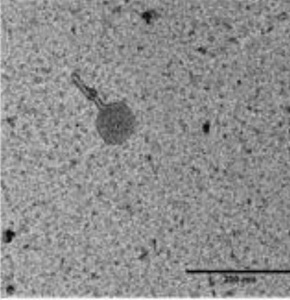 | *Sugarlandvirus* | 109 kb | 36 |
| vB_KpM_SoFaint (Phage 70) | 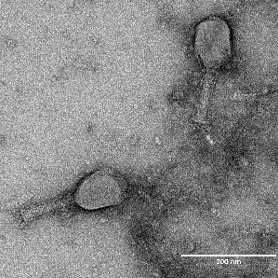 | *Slopekvirus* | 176 kb | 36 |

| **sample_name** | **t0** | **t1** | **t0_idx** | **t1_idx** | **mumax** | **mumax_std** | **dt** | **dt_std** | **doublings** | **doublings_log** | **doublings_log_std** | **yield** | **error** | **smoothing_window** |
| --- | --- | --- | --- | --- | --- | --- | --- | --- | --- | --- | --- | --- | --- | --- |
| TOP52_1 | 20.37 | 158.87 | 4 | 31 | 0.020 | 0.001 | 35.260 | 1.045 | 6.830 | 4.699 | 0 | 1.802 | 0.946 | 10 |
| TOP52_2 | 20.37 | 158.87 | 4 | 31 | 0.020 | 0.001 | 35.063 | 1.696 | 7.620 | 5.224 | 0 | 1.770 | 0.862 | 10 |
| TOP52_3 | 20.37 | 158.87 | 4 | 31 | 0.020 | 0.001 | 34.931 | 1.254 | 6.998 | 4.781 | 0 | 1.798 | 0.908 | 10 |
| 191_1 | 25.43 | 158.87 | 5 | 31 | 0.022 | 0.001 | 31.603 | 1.196 | 8.402 | 5.576 | 0 | 1.850 | 0.876 | 10 |
| 191_2 | 25.43 | 158.87 | 5 | 31 | 0.022 | 0.001 | 32.164 | 1.068 | 7.384 | 4.901 | 0 | 1.881 | 0.927 | 10 |
| 191_3 | 25.43 | 158.87 | 5 | 31 | 0.022 | 0.001 | 32.150 | 0.916 | 8.242 | 5.643 | 0 | 1.897 | 0.901 | 10 |
| 197_1 | 25.43 | 163.93 | 5 | 32 | 0.021 | 0.001 | 33.242 | 1.066 | 7.536 | 5.202 | 0 | 1.831 | 0.906 | 10 |
| 197_2 | 25.43 | 163.93 | 5 | 32 | 0.021 | 0.001 | 33.226 | 1.191 | 7.619 | 5.252 | 0 | 1.822 | 0.894 | 10 |
| 197_3 | 25.43 | 163.93 | 5 | 32 | 0.022 | 0.001 | 31.641 | 0.921 | 7.618 | 5.338 | 0 | 1.860 | 0.918 | 10 |
| 200_1 | 25.43 | 158.87 | 5 | 31 | 0.022 | 0.001 | 32.036 | 1.136 | 7.500 | 5.057 | 0 | 1.822 | 0.911 | 10 |
| 200_2 | 25.43 | 158.87 | 5 | 31 | 0.022 | 0.001 | 32.324 | 0.960 | 7.436 | 4.981 | 0 | 1.882 | 0.932 | 10 |
| 200_3 | 25.43 | 158.87 | 5 | 31 | 0.021 | 0.001 | 31.896 | 0.885 | 7.466 | 5.051 | 0 | 1.886 | 0.936 | 10 |

Supplementary table 2. Growth rate (Mu^max^) and doubling time data from *Klebsiella* Top52 and recovered isolates 191, 197, 200 from Day 7 mice. No significant difference on both growth rate and doubling time from Original Top52 strain and recovered isolates from mice (p<0.05).
